# Genomes of three Sympatric Species of an Adaptive Radiation: *Cyprinodon variegatus, C. desquamator,* and *C. brontotheroides*

**DOI:** 10.64898/2026.10.13.758613

**Authors:** Adam Gomez, Megan A. Barela Hudgell, Evan Lavelle, Joann Mudge, Rhiannon West

## Abstract

Speciation is a central process in evolutionary biology that involves the formation of reproductive barriers between diverging populations. This process can occur through multiple ecological, behavioral, and genetic mechanisms that ultimately lead to changes within a population leading to the formation of distinct species. The relatively young adaptive radiation of *Cyprinodon* pupfishes in the lakes of San Salvador Island provides a powerful system to study the drivers of speciation in species that exist both sympatrically and allopatrically. The three species; *C. variegatus*, *C. desquamator*, and *C. brontotheroides,* can be found in many lakes on the island, exhibit distinct morphological adaptations relating to feeding, and, when in sympatry, the three species have overlapping habitats and breeding grounds. Species boundaries are maintained by asymmetric mate choice by both males and females, which reinforce extreme fitness costs of hybridization. Here we present high-quality annotated reference genomes for these three species generated using PacBio hifi sequencing. The assemblies resulted in a genome size of approximately 1.2 Gb for each species with most sequences scaffolding into 24 chromosomes with BUSCO analysis indicating a >99% level of completion. The annotation of these three genomes generated over 21,500 protein coding genes with BUSCO analysis indicating a >86 completion. These genomes provide a key resource for studying the genomic architecture of ecological specialization, reproductive isolation in sympatry, and rapid speciation and will facilitate future comparative and population genomic studies on adaptive radiations.

## Introduction

Speciation, a core process in evolutionary biology, involves the formation of reproductive barriers between diverging populations (Coyne & Orr, 2004; Nosil, 2012). Contemporary genomic approaches offer unprecedented resolution for characterizing speciation dynamics and elucidating the genetic and evolutionary mechanisms that drive reproductive isolation (Campbell et al., 2018; Nosil, 2012; Seehausen et al., 2014). The *Cyprinodon* radiation on San Salvador Island, The Bahamas, provides an emerging model system for investigating vertebrate diversification, offering insights into the interactions among behavior, ecology, and genomics.

The *Cyprinodon* radiation in the lakes of San Salvador Island represents a young adaptive radiation comprising three species descended from a recent, common ancestor (Martin, 2016). The species (*C. variegatus*, *C. desquamator*, and *C. brontotheroides*) exhibit distinct morphological adaptations, particularly in male nuptial coloration (Kodric-Brown & West, 2014; West & Kodric-Brown, 2015) and trophic morphology, which correspond to specialized dietary niches (Holtmeier, 2001; E. S. Lencer et al., 2016, 2017; Martin, 2016; Martin & Feinstein, 2014; Martin & Wainwright, 2013; McLean & Lonzarich, 2017). They are composed of a generalist omnivore, *Cyprinodon variegatus*, a scale-eater, *C. desquamator*, and a durophage, *C. brontotheroides*. Genomic analyses reveal that selection initially targeted genes associated with feeding behavior, preceding morphological adaptations in the radiation (Richards et al., 2021). While the ancestral-type species, *C. variegatus* is widely distributed across most lakes, the two other species occur sympatrically with the *C. variegatus* in some lakes (Richards & Martin, 2021). In these sympatric populations, the three species share overlapping habitats and breeding grounds, with species boundaries maintained through mate choice (Kodric-Brown & West, 2014; West & Kodric-Brown, 2015) that imposes strong selection against hybridization (Martin & Wainwright, 2013). The unresolved mechanisms driving species distribution patterns across lakes, combined with recent advances in both methodological approaches and theoretical frameworks in speciation research (Peichel et al., 2024), present a compelling opportunity to elucidate the genomic architecture underlying rapid vertebrate diversification in this system.

Emerging evidence indicates that recently diverged sympatric species often exhibit extensive and highly heterogeneous genomic differentiation (Seehausen et al., 2014). Comparative genomic analyses have identified structural variants, such as chromosomal inversions and gene duplications, as potential contributors to the speciation process (Gompert et al., 2025). The annotated genome assemblies presented in this study provide a critical foundation for investigating the role of such genomic features in driving and maintaining species divergence. These high-quality assemblies will enable comparative analyses both within the San Salvadoran radiation and with related species worldwide. Given the large genome size (>1 Gb) and complexity of these diploid species, we employed high-molecular-weight DNA extraction and PacBio HiFi sequencing to ensure accurate assembly of complex regions (Bronner et al., 2025). The resulting assemblies, complemented by available transcriptome data from related species (Jones & Griffitt, 2022; E. Lencer & McCune, 2020; McGirr & Martin, 2019), provide a robust foundation for investigating the genomic basis of speciation in this system (Larsson et al., 2021).

## Methods

### Sample Collection

Four male specimens of *C. desquamator*, one male of *C. brontotheroides*, and one male of *C. variegatus* (whole fish, mean body mass 2.5 g; total length 35 mm), which were collected from San Salvador (Figure 1A) and maintained in aquaria (∼35ppt, room temperature of 24C, ∼800L stock tanks) at Northern New Mexico College, were selected for sequencing. Specimens were euthanized using MS-222 at Northern New Mexico College. Following euthanasia, gut tissue was removed from the *C. desquamator* specimens but maintained in the *C. brontotheroides* and *C. variegatus.* Specimens were immediately flash-frozen in liquid nitrogen in individual 2 mL microcentrifuge tubes. For transport, samples were placed in protective 5 mL Falcon tubes, packed with dry ice, and shipped overnight to Histogenetics (Westbury, NY) for high-molecular-weight DNA extraction and sequencing.

**Figure 1.**
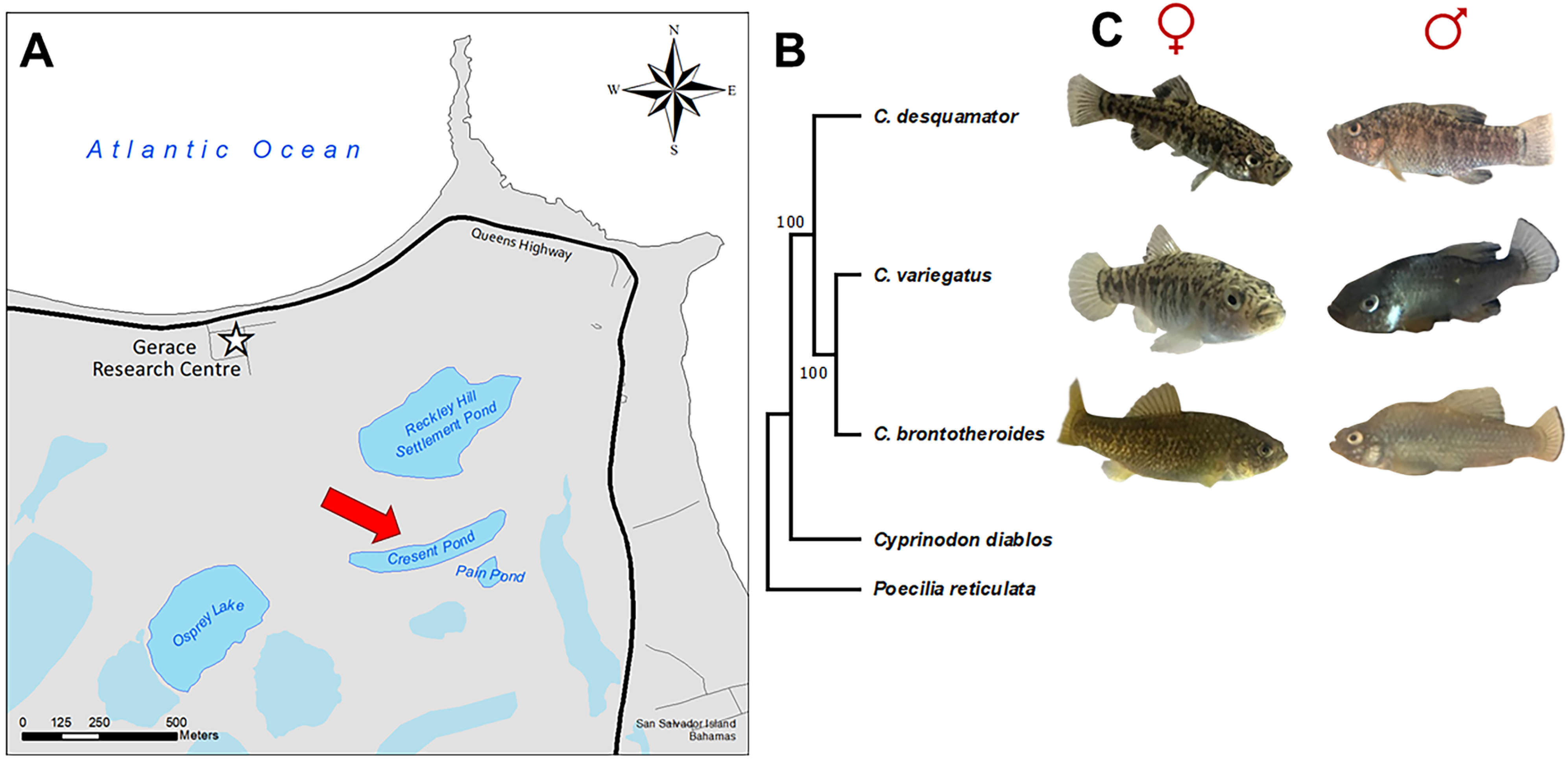
Map of the collection site, phylogeny, and images of the three *Cyprinodon* species whose genomes were assembled. A) Map generated inArcmap 9.2 (Esri, 2006) using data from The San Salvador Island GIS database, compiled by (Robinson & Davis, 1999). Grey indicated the San Salvador landmass while white indicated the Atlantic Ocean. Inland lakes are indicated in blue with Cresent Pond (the collection sight of all three species) being indicated by a red arrow. B) RAxML-HPC2 Phylogenetic tree generated from BUSCO genes of the three study species along with *C. diablos* and *Poecilla reticulata* (outgroup). Bootstrap values were set for 1000 iterations with a bootstrap seed and are indicated to the left of the branch points and the species names are indicated to the right. C) Photographs of the three species of *Cyprinodon* – female (left) and male (right) – species used in this study.

### DNA Extraction, and Sequencing

Genomic DNA was extracted from whole-body sample following the removal of skin from the flash-frozen tissue, 25 mg of the sample was ground using a rotor-stator homogenizer. The tissue was then lysed, and HMW gDNA was purified using the silica-coated magnetic disk (Nanobind tissue kit, PacBio, 102-208-000). To generate long, high-accuracy HiFi reads, the purified gDNA was sheared to a mode size of 18 kb. A second extraction protocol was performed using an equivalent mass of tissue (25 mg) with modified sample preparation (removal of both fins and skin prior to extraction and an optimized tissue homogenization procedure) due to initial yields exhibiting pigmentation and a modal fragment size distribution of ap proximately 10 kb, as determined by gel electrophoresis. This modification eliminated DNA pigmentation however the modal fragment size showed only marginal improvement and remained below the recommended 15-18 kb target size distribution required for optimal Pacific Biosciences High Fidelity (PacBio HiFi) sequencing (Bronner et al., 2024). PacBio HiFi sequencing was completed at Histogenetics (Ossining, NY).

### Genome assembly

PacBio HiFi reads were assessed with FastQC vs 0.11.9 (Andrews, 2010) and assembled using hifiasm v 0.18.5-r499 (Cheng et al., 2021) with default parameters to obtain primary contigs. Assembly contigs were ordered and oriented to create pseudomolecules using *C. diabolis* as a scaffold. Contigs were aligned to *C. diabolis* using minimap2 v2.26-r1175 allowing up to 20% sequence divergence and scaffolds generated by placing 100 Ns between adjacent contigs.

Custom scripts were used to assess assembly statistics. The K-mer Analysis Toolkit (KAT) (Mapleson et al., 2017), which generates k-mer frequency spectra from sequencing reads and compares them against the final assembly, was used to evaluate completeness and identify potential issues such as duplications or missing regions. To further assess completeness, we assessed the percentage of expected Universal Single-Copy Orthologs found in the assemblies using BUSCO v2 (Seppey et al., 2019; Simã et al., 2015; Waterhouse et al., 2018). BUSCO results were generated for all three *Cyprinodon* genomes assembled here along with *Cyprinodon diablos* (GCA_030533445.1) and *Poecilia reticulata* (GCF_000633615.1) using BUSCO 6.0.0. and the Actinopterygii_odb10 lineage.

In addition, comparative visualizations including whole genome dotplots and circos plots were generated (see below). Large scale translocations were checked by visualizing alignments of PacBio HiFi genomic reads to the assemblies (generated using minimap2) in the Integrative Genomics Viewer (IGV) (Thorvaldsdóttir et al., 2013). If reads did not span the junction, the translocation was broken.

### Genome annotation

RNA-Seq data from each species were incorporated to support genome annotation. These include data from BioProjects PRJNA391309 (craniofacial divergence studies in all three species), PRJNA305422 (jaw morphology research), PRJNA508801 (developmental responses to oil and hypoxia exposure), and PRJNA702804 (larval responses to environmental stressors) which encompass diverse tissues including craniofacial structures, head, whole body, whole-embryo, and whole-larvae specimens.

A reference-based assembly was performed as follows. All RNA-Seq data were processed using TrimGalore 0.6.10 (which incorporates Cutadapt and FastQC) for quality control and adapter trimming. Transcriptome reads from each species were aligned to their respective genome assembly using HISAT2 version 2.2 (Kim et al., 2019) and the resulting sam file converted to bam and sorted using samtools version 1.21 (Danecek et al., 2021). The assembly was performed using StringTie v2.2.3 (M. Pertea et al., 2015) and the resulting GTF was converted to GFF3 using gffread v0.12.7 (G. Pertea & Pertea, 2020).

Gene annotations were predicted using MAKER v 3.01.03 (Campbell et al. 2014) following the strategy in (Card et al., 2019). Briefly, RepeatModeler (Smit & Hubley, 2018), which employs computational tools (RECON and RepeatScout) to identify and characterize repeat families - groups of evolutionarily related repetitive sequences derived from common ancestral elements, was used to identify and categorize repeats in the genomes. Then, Repeatmasker version 4.1.5 (Smit et al., 2015) was used to mask the genomes in preparation for gene annotation. The complex repeats were pulled out for use in MAKER and converted into GFF3 format. Two rounds of MAKER were run. The first MAKER round was run passing in the StringTie transcriptome assembly GFF3 file and the Repeatmasker file GFF3 file. Also, existing proteome models from *C. diabolis* (GCA_030533445.1) were used. A second MAKER round was run to improve the gene calls using ESTs, proteins, and repeats identified in the first MAKER run. In addition, Augustus’s zebrafish model and SNAP v.2006-07-28 (Korf, 2004) model trained using high confidence gene models (AED of 0.25 or better and a length of 50 or more amino acids) from *C. variegatus’s* first maker round were used. Assembly completeness was assessed on annotated proteins using BUSCO 6.0.0.

### Mitochondria assembly

The mitochondrial genomes from each fish were identified by first creating a blastable (blastn v. 2.16.0) database for each fish using the contig assembly and then blasting against each database with the *Cyprinodon macularius* mitochondrion sequence (KM985373.1) (Camarena-Rosales et al., 2016). The identified sequences were then analyzed and annotated using MitoAnnotator on MitoFish (Iwasaki et al., 2013; Sato et al., 2018; Zhu et al., 2023).

### Phylogenetic tree

Single-copy BUSCO sequences (see above) were filtered and concatenated to generate a single sequence for each species following workflow outlined in (McMurtrey, 2023). Concatenated sequences were aligned using MAFFT v7.526 (Katoh et al., 2002) with default parameters. A phylogenetic tree was constructed with CIPRES: science gateway (M. A. Miller et al., 2010) using RAxML-HPC v.8 on ACCESS (Stamatakis, 2014) with Bootstrap values set for 1000 iterations with a random bootstrap seed and *Poecilia reticulata* set as an outgroup.

### Comparative visualization

Comparative visualizations were generated using whole genome dotplots as well as circos plots. Dotplots were generated by aligning genome assemblies to *C. diabolis* and to each other using minimap2 (asm20 preset). Plots were generated using DotPlotly (Poorten, 2017) using a minimum alignment length of 20 kb and a minimum aggregate alignment length of 1 Mb.

Circos v 0.69-9 (Krzywinski et al., 2009) was used to visualize alignments between the *Cyprinodon diablos* genome and the contigs/scaffolds of the three Bahamian *Cyprinodon* species. FASTA files were generated using seqkit split2 v2.1.0 (Shen et al., 2016, 2024) with one set containing the largest 40 assembled sequences from the initial contigs assemblies and the second set containing the largest 24 assembled sequences from the scaffold assemblies. These two sets were aligned against the *Cyprinodon diablos* chromosomes using minimap2 version 2.30-r1287 (Li, 2018, 2021) allowing for 10% sequence divergence. Alignments greater than 500 kbp in length were plotted per Circos instructions.

## Results and Discussion

### Genome Assembly and Quality Assessment

High-quality genome assemblies were generated for three closely related San Salvadorian *Cyprinodon* species (Figure 1 B&C, accession numbers: JCEVOQ000000000, JCEVOR000000000, and JCEVOS000000000). The *C. desquamator* genome was assembled from four male specimens which generated 28Gbp for a total coverage of ∼23X assuming a genome size of 1.2 Gb which was assembled into 694 contigs with a total assembly size of 1.22 Gb. The *C. variegatus* was assembled from a single male specimen which generated a total of 32Gbp for a total coverage of ∼27X which was assembled into 1,139 contigs totaling 1.27 Gb.

*C. brontotheroides* was also generated from a single individual yielding 36Gbp for a total coverage of ∼30X which generated 1,121 contigs with a total length of 1.26 Gb. All three assemblies demonstrated high contiguity, with N50 values ranging from 26.36 Mb to 28.81 Mb and consistent L50 values of 17 across all species (Table 1). The contig assembly was scaffolded against the *C. diablos* genome (GCA_030533445.1) to create pseudomolecules. The scaffold and contig metrics in all assemblies indicate high contiguity for each genome. Alignment of the largest 40 contigs against *Cyprinodon diablos* indicated that many of the chromosomes were captured by a single contig (*C. desquamator*: 11 chromosomes, *C. brontotheroides*: 9 chromosomes, *C. variegatus* 8 chromosomes) with the remaining chromosomes split into only a few contigs (Supplemental Figure 1).

**Table 1:** Comparative contig assembly statistics for three *Cyprinodon* species.

|  | *C. desquamator* | *C. variegatus* | *C. brontotheroides* |
| --- | --- | --- | --- |
| Number of Contigs | 694 | 1,139 | 1,121 |
| Total Length (Gb) | 1.22 | 1.27 | 1.26 |
| Contig N50 (Mb) | 28.81 | 26.36 | 27.86 |
| Contig L50 | 17 | 17 | 17 |
| Largest Contig (Mb) | 50.28 | 79.81 | 47.3 |
| Smallest contig (kb) | 14.2 | 11 | 12.43 |

The number of scaffolds was nearly halved with *C. desquamator* having 370 scaffolds, *C. variegatus* having 652 scaffolds, and *C. brontotheroides* having 587 scaffolds (Table 2). Of these scaffolds the largest 24 were identified as likely pseudochromosomes with the largest 24 containing ∼95% (*C. desquamator)*, ∼94% (*C. variegatus)*, and ∼93% (*C. brontotheroides)* of the incorporated base pairs. N50 values doubled in size with the average now at 50Mb. The L50 value also increased to 12, reflecting the capture of full-length chromosomes. The largest contig of *C. variegatus* (79.81 Mb) was identified as being a concatenation of two chromosomes (Cv12 and Cv20) that was not supported by the reads. It was, therefore, split in the generation of the final pseudomolecules (Supplemental Figure 1B).

**Table 2:** Comparative pseudomolecule assembly statistics for three *Cyprinodon* species.

|  | *C. desquamator* | *C. variegatus* | *C. brontotheroides* |
| --- | --- | --- | --- |
| Number of Scaffolds | 370 | 652 | 587 |
| Total Length (Gb) | 1.22 | 1.27 | 1.26 |
| Scaffold N50 (Mb) | 49.14 | 50.41 | 50.45 |
| Scaffold L50 | 12 | 12 | 12 |
| Largest Scaffold (Mb) | 58.84 | 60.63 | 61.44 |
| #N's | 2.68 | 3.84 | 4.23 |

To assess genome completeness, we evaluated each assembly using BUSCO (Simã et al., 2015) with the Actinopterygii lineage dataset (Figure 2). Each genome was nearly complete, capturing >99% of conserved single-copy orthologs. *C. desquamator* and *C. brontotheroides* both achieved 99.3% completeness, while *C. variegatus* showed 99.1% completeness. Interestingly, we observed varying levels of gene duplication among the species, with *C. brontotheroides* showing the highest proportion of duplicated single-copy orthologs (3.6%), followed by *C. variegatus* (2.5%) and *C. desquamator* (1.6%). All three genomes had very few missing BUSCOs (0.2-0.3%) and similarly low levels of fragmented genes (0.5-0.6%), further supporting the high quality of these assemblies.

**Figure 2.**
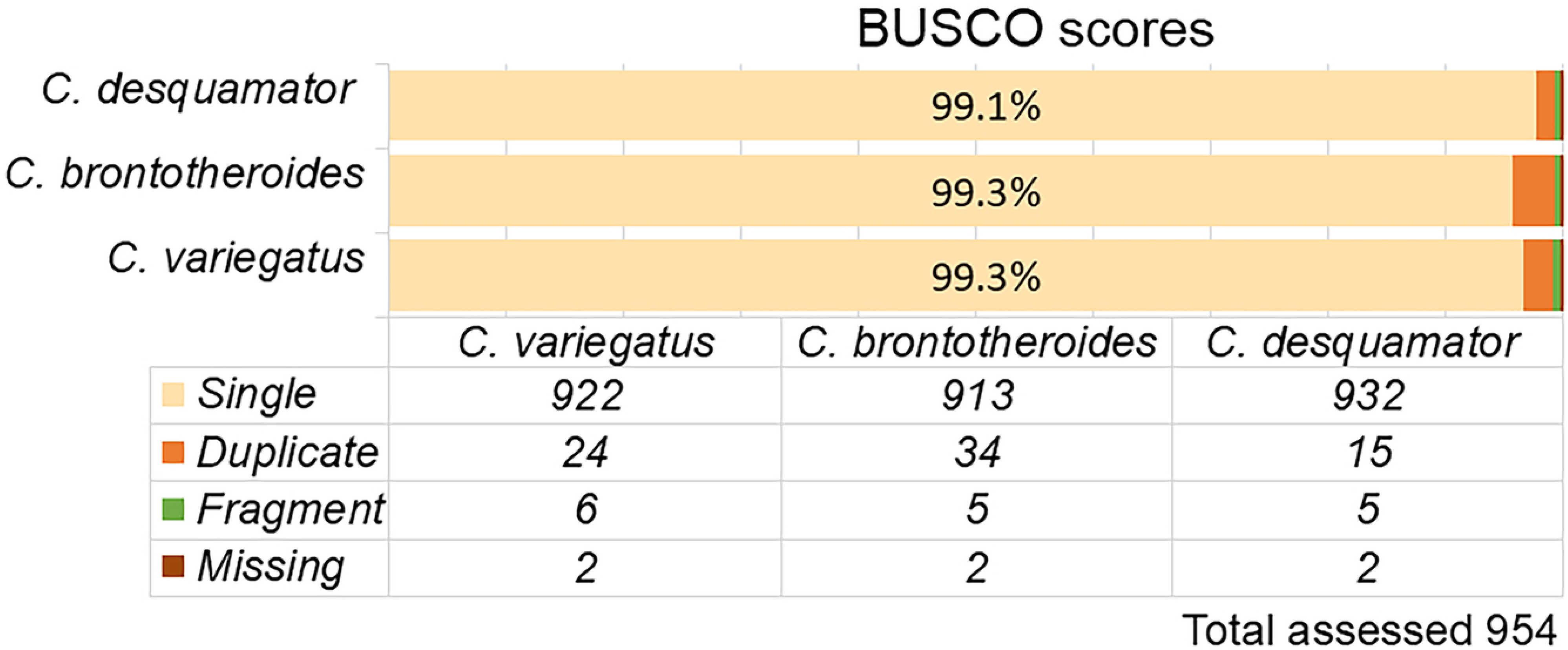
Busco analysis data for the three *Cyprinodon* genomes. BUSCO scores were calculated using 954 genes from the Actinopterygii lineage dataset. The complete dataset is presented below the bar chart.

To assess genome characteristics and potential assembly challenges, we performed k-mer analysis. The k-mer comparison plots (Supplementary Figure 2) show the distribution of k-mers in the reads (black) versus their presence in the final assembly (red, indicating single copy regions; purple, representing potential duplications). The high proportion of k-mers occurring exactly once in both reads and assembly (red peak) indicates good coverage and assembly completeness, while the minimal black regions suggest low levels of unassembled content.

#### Comparative Genomic Analysis

*C. desquamator, C. variegatus*, and *C. brontotheroides* genomes were scaffolded and compared to the *C. diablos* chromosomes as the closest related *Cyprinodon* with a chromosome level assembly. Circos Comparison to *C. diablos* indicated a high level of similarity in chromosome layout across these species with very few small translocations being evident across chromosomes (Figure 3A-C). Because of this the chromosome numbers from the *C. diablos* assembly were used to number the chromosomes in the three *Cyprinodon* genome assemblies presented here. While these three genomes were scaffolded using *C. diabolis*, which could introduce some bias, chromosome squashes across the *Cyprinodon* order all indicate a 2N of 48 (n=24) which gives confidence to the assembly of these genomes into 24 chromosomal units (Baksi & Means, 1988; R. Miller & Echelle, 1975; Nirchio et al., 2003; Turner & Liu, 1977). In general, the chromosome sizes in the three assembled *Cyprinodon* chromosomes are larger than those seen in *C. diablos* with chromosomes 14 and 24 showing a significant increase in size (Table 3).

**Figure 3.**
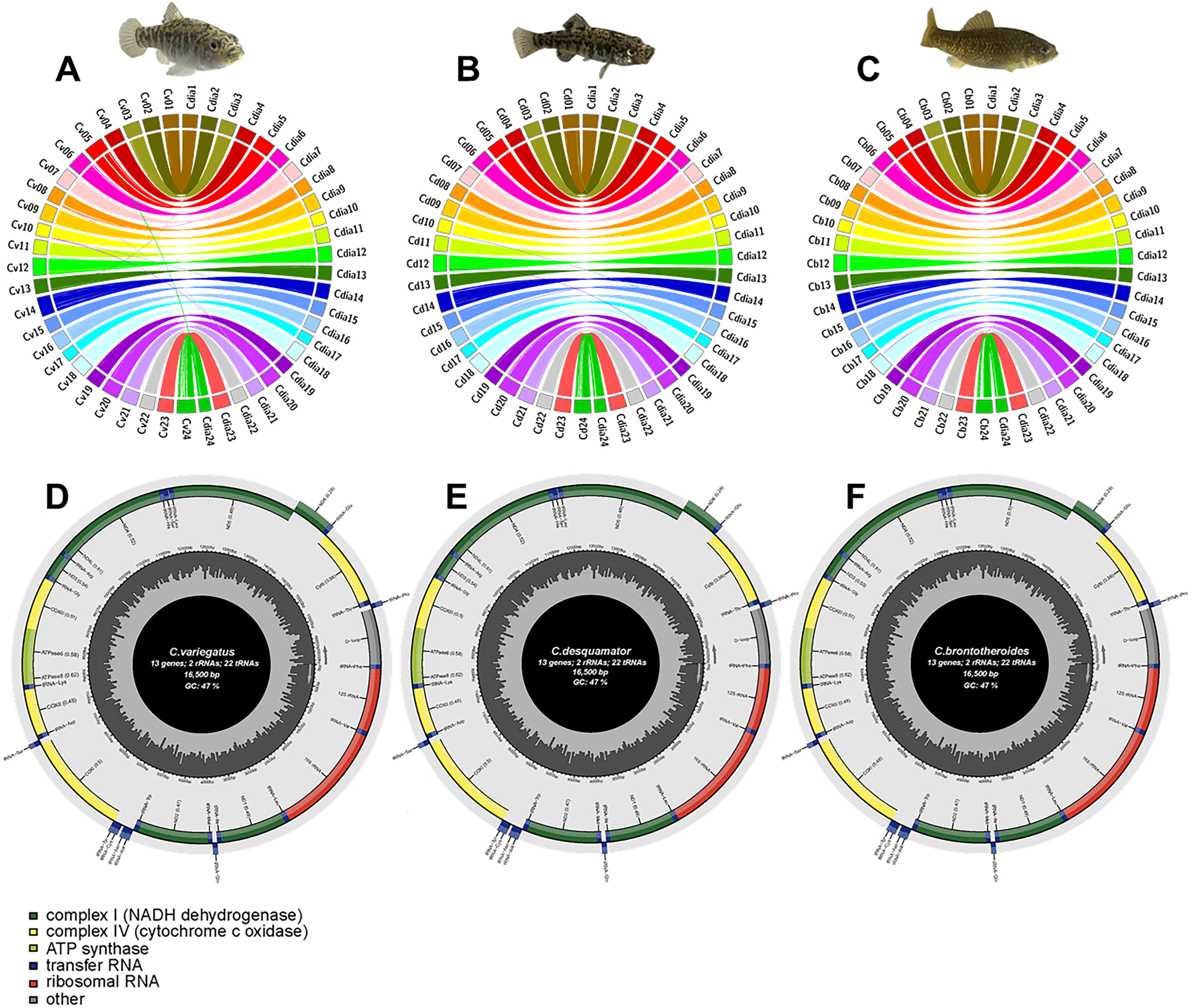
*C. desquamator, C. brontotheroides and C. variegatus* genome and mitochondrial analysis. Scaffolds and chromosomes are represented by solid-colored blocks. Colored ribbons within the circle represent regions of matching sequence greater than 500 kbp. “Cdia” labels indicate *C. diablos* chromosomes across all three circos plots. A) Comparison of *C. variegatus* contigs (labeled Cv) against *C. diablos* chromosomes. B) Comparison of *C. desquamator* scaffolds (labeled Cd) against *C. diablos* chromosomes. C) Comparison of *C. brontotheroides* contigs (labeled Cb) against *C. diablos* chromosomes. D) annotated mitochondrial genome map for *C. variegatus* showing 13 protein coding genes, 2 ribosomal RNAs, 22 transfer RNAs and the control region (Dloop). E) Annotated mitochondrial genome map for *C. desquamator* F) Annotated mitochondrial genome map for *C. brontotheroides*.

**Table 3:** Chromosome size comparison across Cyprinodons.

| Table 3: Chromosome size comparison across *Cyprinodons* |  |  |  |  |
| --- | --- | --- | --- | --- |
|  | *C. diablos* | *C. variegatus* | *C. desquamator* | *C. brontotheroides* |
| Chromosome | Size (bp) |  |  |  |
| 1 | 48,130,825 | 50,279,010 | 52,210,586 | 49,342,378 |
| 2 | 50,011,971 | 50,470,613 | 49,514,870 | 50,710,611 |
| 3 | 49,045,530 | 50,876,134 | 47,456,562 | 49,599,294 |
| 4 | 53,884,543 | 52,577,974 | 52,104,337 | 52,019,878 |
| 5 | 55,109,245 | 60,631,312 | 51,480,684 | 52,823,892 |
| 6 | 48,811,528 | 56,765,708 | 52,919,135 | 53,641,129 |
| 7 | 50,015,530 | 53,249,464 | 52,218,926 | 52,073,639 |
| 8 | 43,920,048 | 47,870,972 | 44,855,259 | 44,994,049 |
| 9 | 49,128,472 | 50,419,012 | 49,141,782 | 50,794,084 |
| 10 | 39,508,766 | 40,308,427 | 42,023,889 | 37,645,509 |
| 11 | 46,303,725 | 45,606,627 | 47,168,322 | 48,313,148 |
| 12 | 50,123,716 | 55,034,608 | 52,809,365 | 53,570,884 |
| 13 | 44,416,577 | 45,035,111 | 44,048,398 | 46,188,723 |
| 14 | 48,018,592 | 60,454,162 | 58,847,056 | 61,444,455 |
| 15 | 42,424,397 | 43,133,479 | 43,033,478 | 41,915,611 |
| 16 | 42,990,238 | 46,709,368 | 46,877,422 | 46,754,001 |
| 17 | 31,953,358 | 34,043,719 | 33,146,809 | 33,546,442 |
| 18 | 49,935,463 | 51,297,870 | 50,995,423 | 54,465,479 |
| 19 | 41,517,714 | 47,870,972 | 44,523,948 | 42,431,543 |
| 20 | 51,780,930 | 53,172,106 | 55,368,452 | 54,709,286 |
| 21 | 45,011,060 | 49,808,126 | 48,397,349 | 49,916,907 |
| 22 | 44,145,131 | 43,843,317 | 43,753,483 | 42,946,567 |
| 23 | 50,711,760 | 49,013,558 | 48,956,113 | 50,456,442 |
| 24 | 35,868,736 | 58,313,722 | 51,945,892 | 50,630,815 |

Additional analysis of the three *Cyprinodon* genomes against one another indicated high levels of conservation across all chromosomes (See Supplemental Figure 3) with dotplots comparing the chromosomes showing a clean diagonal line with very few breaks and inversions detected. A more in-depth analysis comparing these three *Cyprinodon* genomes is pending.

#### Mitochondrial analysis

There is high conservation of the mitochondria chromosome across all three Bahamian pupfish species. All are exactly 16500bp in length. All have 13 genes identified, 2 rRNAs and 22 tRNAs with GC content at 47%, reflecting their shared history (Figure 3 D – F). There is a total of 22 bp differences between *C. desquamator* versus *C. variegatus* and 24 bp differences versus *C. brontotheroides* while there are 18 bp differences between *C. variegatus* and *C. brontotheroides.* These differences are dispersed throughout the length of the mitochondrial genome indicating that if selection is driving these changes, it is acting at multiple locations. Given that these data are generated from a limited number of individuals, it is premature to use these mitochondrial sequences to distinguish these species. Instead, morphological identification is currently the best way to differentiate these species.

### Genome Annotation

#### Repeat Annotation

RepeatModeler identified between 43.89 and 45.14% of interspersed repeats within these three *Cyprinodon* genomes (Table 4, Figure 4). The largest category of these repeats were unclassified (21.98-23.25%) with the next largest category being DNA transposons (11.10-11.44%), in particular the hobo-activator superfamily. Overall, the identified repeats were similar across all three species with little variation with the exception of the 0.00% identification of Penelope superfamily LINE retroelements in *C. desquamator* while the other two species showed a small (541-562bp/0.01-0.02%) number of elements containing this superfamily. On the other hand, *C. desquamator* seemed to have an increased number of Ty1/Copia LTR retroelements (0.27%) compared to the other species, both of which only have 0.03%. The last point of note is the reduction in rolling-circles in the *C. brontotheroides* (0.47%) compared to the other two species (1.10-1.41%).

**Figure 4.**
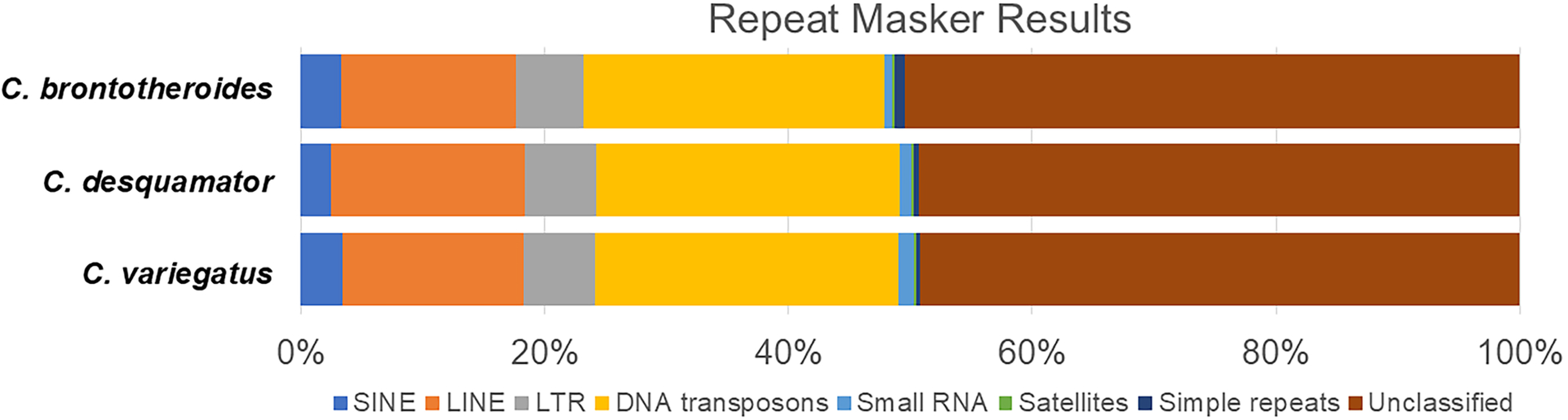
Repeat Masker data for the three *Cyprinodon* genomes. Bar chart comparing the percentage of each repeat order found within each *Cyprinodon* genome as calculated by Repeat Masker. See Table 3 for a more detailed breakdown of results

**Table 4:**
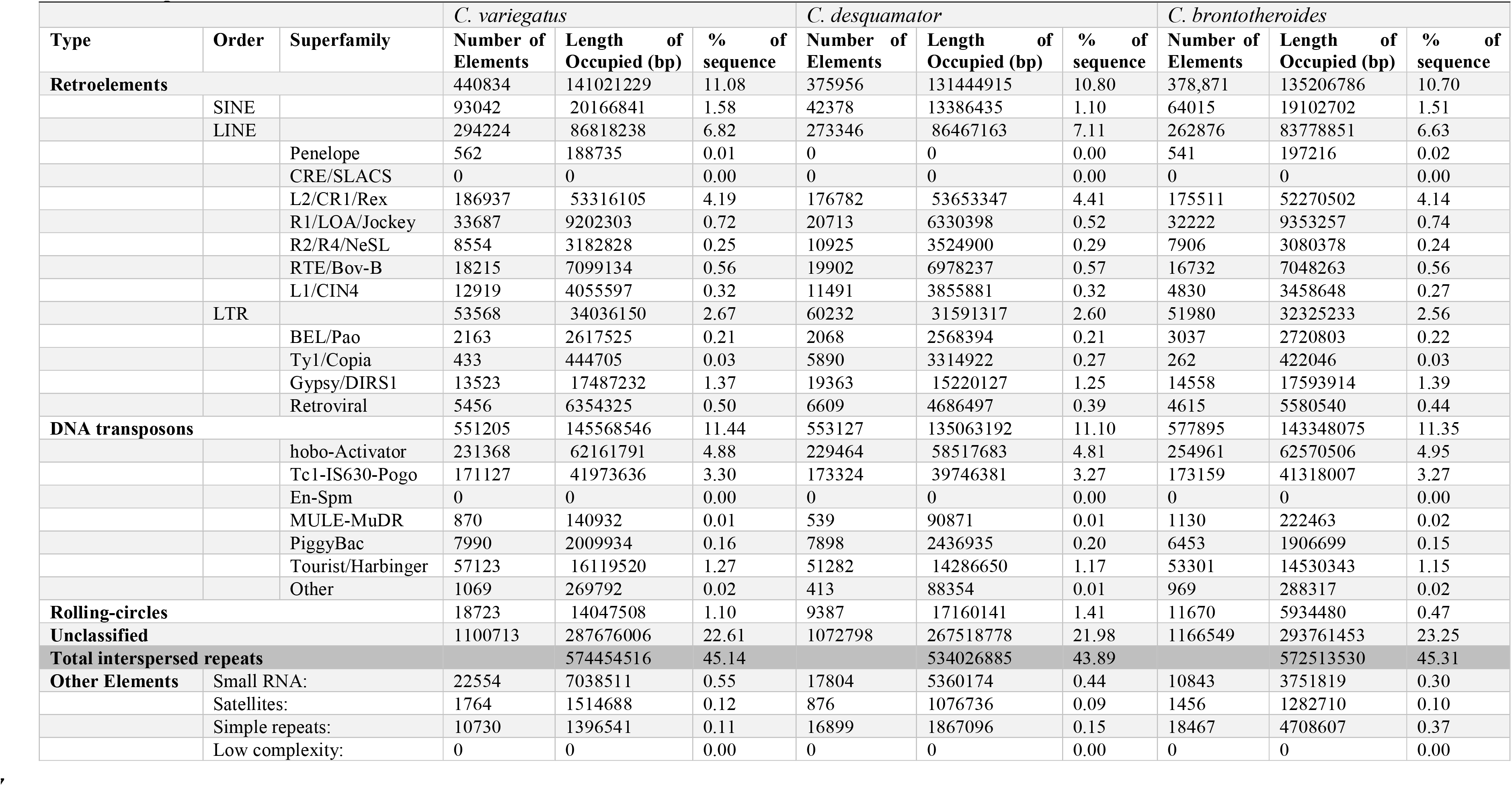
Repeat data.

#### RNA-Seq Evidence

Conspecific RNA-seq datasets were used to provide evidence for gene annotation for each species. *C. variegatus* had the largest dataset available with 119 paired-end Illumina samples from five bioprojects, which encompassed a diverse tissue set including craniofacial structures, head, whole body, whole-embryo, and whole-larvae specimens. The other two species each had 21 samples. These samples included whole-embryo, whole-larvae, whole body, and craniofacial tissues. All three species had a comprehensive transcriptomic dataset across different developmental stages and anatomical regions, enabling robust gene prediction and annotation, particularly for genes involved in morphological development, environmental adaptation, and stress response pathways. The diverse tissue sampling enables detailed investigation of the genetic basis of trophic specialization in this species, including for genes involved in craniofacial development and morphological adaptation. RNA-seq read statistics, including information about trimming, is available for each species in Supplemental Table 1. *C. brontotheroides* and *C. desquamator* each had similar number of reads (671M and 652M, respectively), while *C. variegatus* had many more at 5219M reads.

Following quality control, reads were aligned to their respective genome assemblies using HISAT2. The alignment statistics for each species are presented in Supplemental Table 2.

#### Gene Annotation

We identified 21,653, 22,235, and 21,594 protein coding genes in the *C. variegatus*, *C. brontotheroides*, and *C. desquamator* genome assemblies, respectively (Table 5). This is likely an underestimate of the number of genes given that BUSCO completeness scores on the proteins indicate that the annotation is missing just under 10% of the single copy orthologues. Nevertheless, many of these are captured (but not annotated) in the assembly given genome-wide BUSCO scores above 99% in each assembly (see above). The number of protein coding genes is comparable to those in other pupfish (*C. tularosa* (GCF_016077235.1): 23,979 and *C. variegatus* from Florida (GCF_000732505.1) 23,277) (Black et al., 2021; E. S. Lencer et al., 2017). The genes are high quality, with more than 93% of the gene calls being high confidence (having an annotation edit distance (AED) score >= 0.5. In addition, the vast majority of the annotated protein-coding genes were captured on the pseudochromosomes with 2% or less occurring on scaffolds not incorporated into the pseudochromosomes.

**Table 5:** Gene Annotations.

|  | *C. variegatus* | *C. brontotheroides* | *C. desquamator* |
| --- | --- | --- | --- |
| Protein-coding Genes | 21,653 | 22,235 | 21,594 |
| Genes on chrs | 21,349 | 21,781 | 21,365 |
| % genes on chrs | 98.6% | 98.0% | 98.9% |
| tRNA genes (pseudo) | 10,236 (2,578) | 9,613 (2,514) | 9,873 (2,510) |
| Complete BUSCOs | 86.2% | 86.1% | 86.2% |
| Missing BUSCOs | 9.6% | 9.6% | 9.5% |
| High confidence gene calls | 93.6% | 93.8% | 94.2% |

## Summary

The pupfishes of Cyprinodon on San Salvador Island, The Bahamas, represent a young vertebrate adaptive radiation that includes three closely related species: the widespread generalist *C. variegatus* and two trophic specialists, the scale-eater *C. desquamator* and the durophage *C. brontotheroides*. In several lakes, these species occur sympatrically, occupying overlapping habitats and breeding sites while maintaining species boundaries primarily through assortative mating and strong selection against hybridization. This system therefore provides a powerful natural framework for examining the genomic basis of reproductive isolation and ecological diversification in sympatry. Here, we present high-quality annotated genome assemblies for the three San Salvador pupfish species. These assemblies were generated using high-molecular-weight DNA and PacBio HiFi sequencing to resolve the large and complex (>1 Gb) genomes characteristic of this clade. Together with existing transcriptomic resources, these genomes establish a comparative framework for investigating the genetic architecture of rapid ecological specialization, the role of structural variation in maintaining species boundaries, and the broader mechanisms underlying sympatric speciation in vertebrates.

## Acknowledgments

We thank A. Dichosa for valuable discussions and ongoing continual support of this work; M. Ritchie and the Friday Lab Chat members for discussion and feedback; V. Benavidez, E. Gonzalez, J. Gutierrez, and J. Madrid for their unflagging assistance in the field and members of the West Pupfish Lab for care and support of fish and each other; and T. Atencio, J. Gallegos, I. Lopez, S. Nandy, C. Pacheco, M. Romero, NNMC Facilities, the NNMC business office, NMSU-INBRE, and T. Dexter and the staff of the Gerace Research Centre for logistical support, without whom none of this would have been possible - science takes a community. Finally, with the deepest gratitude, we thank the people of the island of San Salvador for welcoming researchers into their community. The work was conducted under the Bahamian permits BEST-AGR/NAT/1:33/2018 and BS-2024-788486 and NNMC IACUC approvals: IACUC-2018-PI-RW, IACUC-2021-PI-RW-R, and IACUC-2024-PI-RW-R.

## Data Access

The genome assemblies and annotations from this study have been submitted to NCBI Genbank under accession numbers JCEVOQ000000000, JCEVOR000000000, and JCEVOS000000000 (release pending). Supplementary File 1 contains supplementary figures and tables.

## Ethics and Approval

This work was approved by the Northern New Mexico College Institutional Animal Care and Use Committee (IACUC), Animal Welfare Assurance D21--01105, USDA Registration NA, under protocol IACUC-2018-pt-Rw-v.2.

