## Supplementary File 1 for "Genomes of three Sympatric Species of an Adaptive Radiation: *Cyprinodon variegatus, C. desquamator,* and *C. brontotheroides*"

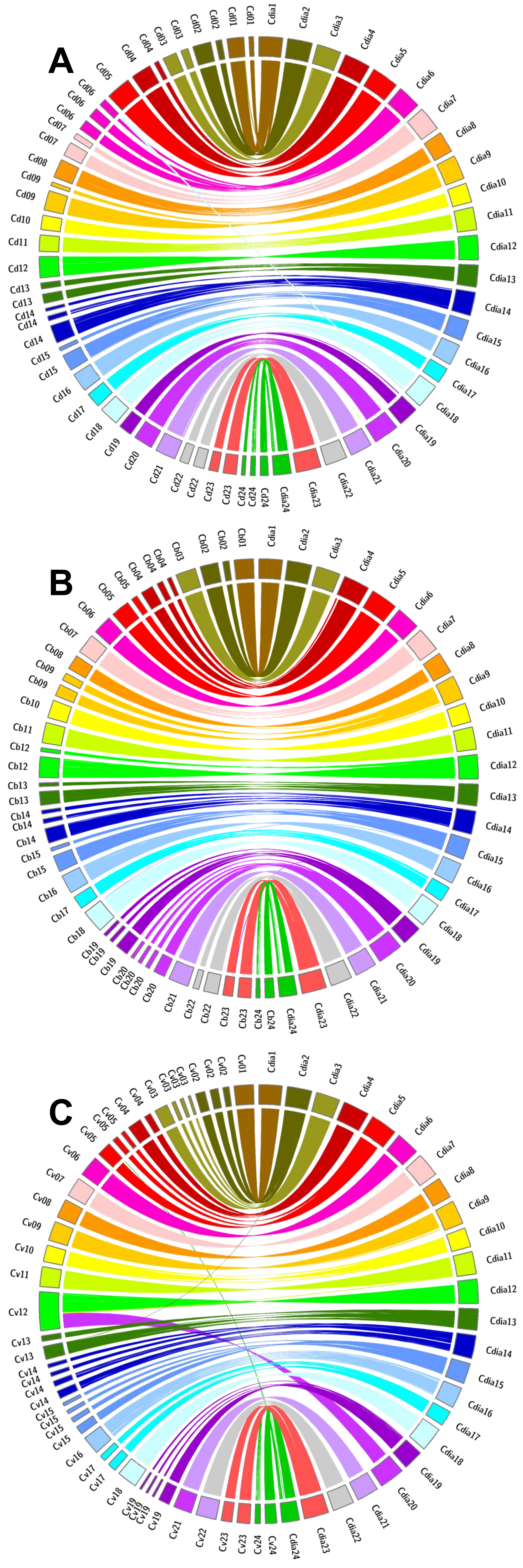
A
Cd01
Cd02
Cd03
Cd04
Cd05
Cd06
Cd07
Cd08
Cd09
Cd10
Cd11
Cd12
Cd13
Cd14
Cd15
Cd16
Cd17
Cd18
Cd19
Cd20
Cd21
Cd22
Cd23
Cd24
Cdia1
Cdia2
Cdia3
Cdia4
Cdia5
Cdia6
Cdia7
Cdia8
Cdia9
Cdia10
Cdia11
Cdia12
Cdia13
Cdia14
Cdia15
Cdia16
Cdia17
Cdia18
Cdia19
Cdia20
Cdia21
Cdia22
Cdia23
Cdia24
B
Cb01
Cb02
Cb03
Cb04
Cb05
Cb06
Cb07
Cb08
Cb09
Cb10
Cb11
Cb12
Cb13
Cb14
Cb15
Cb16
Cb17
Cb18
Cb19
Cb20
Cb21
Cb22
Cb23
Cb24
C
Cv01
Cv02
Cv03
Cv04
Cv05
Cv06
Cv07
Cv08
Cv09
Cv10
Cv11
Cv12
Cv13
Cv14
Cv15
Cv16
Cv17
Cv18
Cv19
Cv21
Cv22
Cv23
Cv24

**Supplementary Figure 1 | Circos plot of the largest 40 contigs from three *Cyprinodon* assemblies compared to *C. diablos* chromosomes.** Contigs and chromosomes are represented by solid-colored blocks. Colored ribbons within the circle represent regions of matching sequence greater than 500 kbp. "Cdia" labels indicate *C. diablos* chromosomes across all three circos plots. A) Comparison of *C. desquamator* contigs (labeled Cd) against *C. diablos* chromosomes. B) Comparison of *C. brontotheroides* contigs (labeled Cb) against *C. diablos* chromosomes. C) Comparison of *C. variegatus* contigs (labeled Cv) against *C. diablos* chromosomes.

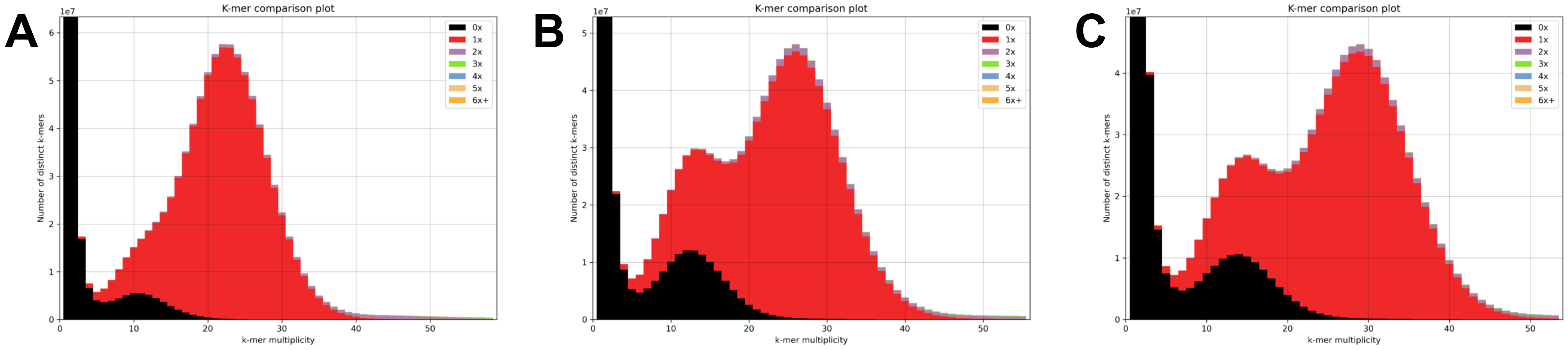
A
B
C
K-mer comparison plot
1e7
Number of distinct k-mers
k-mer multiplicity
0x
1x
2x
3x
4x
5x
6x+

**Supplementary Figure 2 | K-mer comparison plots generated by the K-mer Analysis Toolkit (KAT) for the three Cyprinodon genomes.** Each plot shows the k-mer frequency spectrum of the sequencing reads (black line) compared to their occurrence in the final assembly. Red coloring indicates k-mers present once in the assembly (properly assembled unique regions), while purple indicates k-mers present multiple times (potential duplications). The x-axis represents k-mer multiplicity in the reads, and the y-axis shows the number of distinct k-mers at each multiplicity.

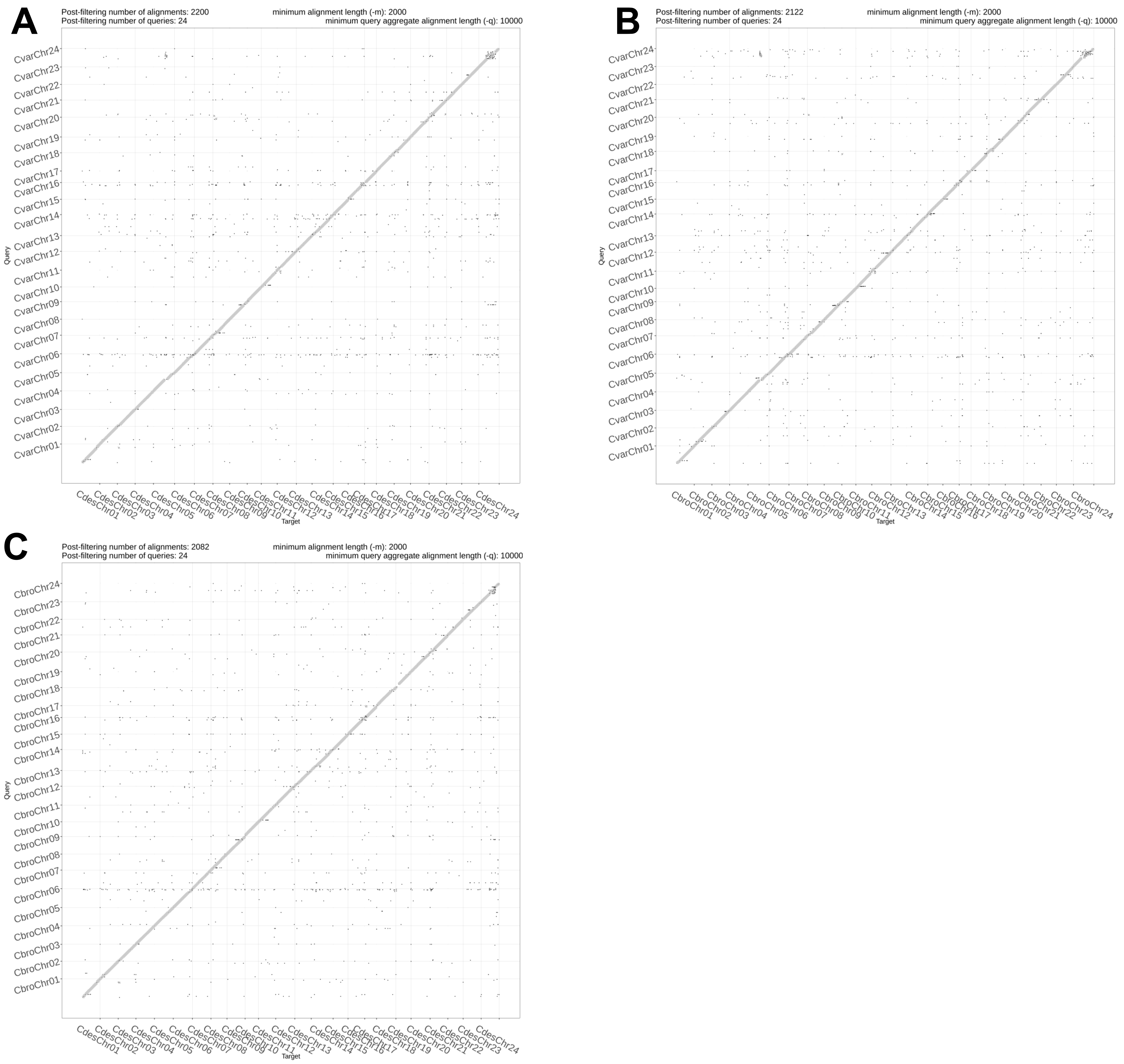
A
Post-filtering number of alignments: 2200
Post-filtering number of queries: 24
minimum alignment length (-m): 2000
minimum query aggregate alignment length (-q): 10000
Query
CvarChr01 CvarChr02 CvarChr03 CvarChr04 CvarChr05 CvarChr06 CvarChr07 CvarChr08 CvarChr09 CvarChr10 CvarChr11 CvarChr12 CvarChr13 CvarChr14 CvarChr15 CvarChr16 CvarChr17 CvarChr18 CvarChr19 CvarChr20 CvarChr21 CvarChr22 CvarChr23 CvarChr24
Target
CdesChr01 CdesChr02 CdesChr03 CdesChr04 CdesChr05 CdesChr06 CdesChr07 CdesChr08 CdesChr09 CdesChr10 CdesChr11 CdesChr12 CdesChr13 CdesChr14 CdesChr15 CdesChr16 CdesChr17 CdesChr18 CdesChr19 CdesChr20 CdesChr21 CdesChr22 CdesChr23 CdesChr24
B
Post-filtering number of alignments: 2122
Post-filtering number of queries: 24
minimum alignment length (-m): 2000
minimum query aggregate alignment length (-q): 10000
Query
CvarChr01 CvarChr02 CvarChr03 CvarChr04 CvarChr05 CvarChr06 CvarChr07 CvarChr08 CvarChr09 CvarChr10 CvarChr11 CvarChr12 CvarChr13 CvarChr14 CvarChr15 CvarChr16 CvarChr17 CvarChr18 CvarChr19 CvarChr20 CvarChr21 CvarChr22 CvarChr23 CvarChr24
Target
CbroChr01 CbroChr02 CbroChr03 CbroChr04 CbroChr05 CbroChr06 CbroChr07 CbroChr08 CbroChr09 CbroChr10 CbroChr11 CbroChr12 CbroChr13 CbroChr14 CbroChr15 CbroChr16 CbroChr17 CbroChr18 CbroChr19 CbroChr20 CbroChr21 CbroChr22 CbroChr23 CbroChr24
C
Post-filtering number of alignments: 2082
Post-filtering number of queries: 24
minimum alignment length (-m): 2000
minimum query aggregate alignment length (-q): 10000
Query
CbroChr01 CbroChr02 CbroChr03 CbroChr04 CbroChr05 CbroChr06 CbroChr07 CbroChr08 CbroChr09 CbroChr10 CbroChr11 CbroChr12 CbroChr13 CbroChr14 CbroChr15 CbroChr16 CbroChr17 CbroChr18 CbroChr19 CbroChr20 CbroChr21 CbroChr22 CbroChr23 CbroChr24
Target
CdesChr01 CdesChr02 CdesChr03 CdesChr04 CdesChr05 CdesChr06 CdesChr07 CdesChr08 CdesChr09 CdesChr10 CdesChr11 CdesChr12 CdesChr13 CdesChr14 CdesChr15 CdesChr16 CdesChr17 CdesChr18 CdesChr19 CdesChr20 CdesChr21 CdesChr22 CdesChr23 CdesChr24

**Supplementary Figure 3 | Pairwise dotplots of all three *Cyprinodon* genome assemblies.** The diagonal lines in each dotplot indicate regions of similar sequence. Breaks in the diagonal indicate mismatches and inversions are characterized by a change in the direction of the diagonal. The dot observed around the dotplot indicate areas of matching sequence outside of the diagonal. A) *C. variegatus* chromosomes (labeled Cvar) against *C. desquamator* chromosomes (Cdes) B) *C. variegatus* chromosomes (Cvar) against *C. brontotheroides* chromosomes (Cbro) C) *C. brontotheroides* chromosomes (Cbro) against *C. desquamator* (Cdes).

Supplemental Table 1: RNA-Seq data processing statistics for the three Cyprinodon species

| | *C. variegatus* | *C. brontotheroides* | *C. desquamator* |
|---|---|---|---|
| **Total Reads** | 5,219,338,104 | 670,675,974 | 652,429,874 |
| **Reads with Adapters** | 2,448,713,780 (46.9%) | 267,009,658 (39.8%) | 242,366,342 (37.1%) |
| **Reads Passing Filters** | 5,219,338,104 (100%) | 670,675,974 (100%) | 652,429,874 (100%) |
| **Total Basepairs** | 697,982,967,666 | 100,790,460,968 | 97,900,285,850 |
| **Quality-trimmed** | 3,540,000,493 (0.5%) | 1,168,224,572 (1.2%) | 1,143,847,505 (1.2%) |
| **Total Written** | 658,790,245,014 (94.4%) | 97,280,423,393 (96.5%) | 95,098,253,785 (97.1%) |
| **Pairs Removed** | 3,521,995 (0.13%) | 867,677 (0.26%) | 787,658 (0.24%) |

Supplemental Table 2: RNA-Seq alignment statistics for the three Cyprinodon species

| | *C. variegatus* | *C. brontotheroides* | *C. desquamator* |
|---|---|---|---|
| **Total Reads** | 5,212,294,114 | 668,940,620 | 650,854,558 |
| **Mapped Reads** | 4,580,979,541 (87.9%) | 568,260,121 (84.9%) | 573,554,069 (88.1%) |
| **Properly Paired** | 4,350,445,942 (95.0%) | 549,380,950 (96.7%) | 555,042,760 (96.8%) |
| **Unmapped Reads** | 631,314,573 (12.1%) | 100,680,499 (15.1%) | 77,300,489 (11.9%) |
| **Bases Mapped** | 575,543,525,436 | 82,820,597,434 | 83,907,219,131 |
| **Pairs Removed** | 3,521,995 (0.13%) | 867,677 (0.26%) | 787,658 (0.24%) |
